# Random Parametric Perturbations of Gene Regulatory Circuit Uncover State Transitions in Cell Cycle

**DOI:** 10.1101/799965

**Authors:** Ataur Katebi, Vivek Kohar, Mingyang Lu

## Abstract

Many biological processes involve precise cellular state transitions controlled by complex gene regulation. Here, we use budding yeast cell cycle as a model system and explore how a gene regulatory circuit encodes essential information of state transitions. We present a generalized random circuit perturbation (RACIPE) method, specifically for circuits containing heterogeneous regulation types, and its usage to analyze both stable steady states and oscillatory states from an ensemble of circuit models with random kinetic parameters. The stable steady states form robust clusters with a circular structure that are associated with cell cycle phases. We show that this circular structure in the clusters is consistent with single cell RNA-seq data. The oscillatory states specify irreversible state transitions along cell cycle progression. Furthermore, we identify possible mechanisms to understand irreversible state transitions from steady states of random models. We expect this approach to be robust and generally applicable to unbiasedly predict dynamical transitions of a gene regulatory circuit.

---

One of the most significant challenges in biology is to elucidate the function and control mechanism of gene regulatory circuits (GRCs) that govern the decision making of biological processes^1,2^. The rapid development of new genomics technology has allowed us to address this question by both experimental and computational systems biology approaches^3–5^. One of the critical components among them is to use mathematical modeling to simulate the dynamical behavior of GRCs^6–8^. A good model not only provides a mechanistic view of the system but also makes new predictions on the effect of a genetic perturbation like the knockdown and overexpression of one or more genes. While mathematical modeling approaches have been successfully applied to analyze gene circuits in select cases^9–13^, they typically suffer from a long-standing problem inherent in almost every systems biology study -- it is difficult to directly measure most circuit kinetic parameters, especially in vivo. Although some parameter values can be learned from published results, many others are often based on educated guesses, therefore, limiting the predictive power of mathematical modeling. The problem is amplified when circuit modeling is applied to large gene networks because such systems typically have many parameters, making conventional methods prone to over-fitting.

To address this issue, we have recently developed a mathematical modeling algorithm, named *ra*ndom *ci*rcuit *pe*rturbation (RACIPE)^14^, which captures the dynamics of a GRC without the need for precise kinetic parameters^15^. Instead of fine-tuning a model with a specific set of parameters, RACIPE generates an ensemble of models based on the associated chemical rate equations with distinct random kinetic parameter sets. The random models can represent the GRC in different signaling states and/or microenvironments. The simulated expressions from the ensemble of models are then subjected to statistical analysis to identify robust features of the GRC. Particularly, we found, in several applications to simple circuit motifs and biological circuits16, that even though the kinetic parameters are largely randomized, the stable steady state solutions from these models form robust clusters of states that can be associated with distinct cellular states. This finding suggests that we can use circuit topology (i.e., the connectivity and the types of regulatory interactions) as the only input and unbiasedly predict the cellular states that are allowed by the circuit. Without the need to refine a particular set of parameters, RACIPE can be extended to simulate a large gene network. In essence, RACIPE uniquely converts the nonlinear dynamics problem to a statistical data analysis problem, an unbiased approach for analyzing large network-systems.

Ever since its development, RACIPE has helped substantially, in several cases^15–18^, to understand the possible gene expression steady states and the roles of genes in the behavior of the GRCs. It remains unclear how efficient the ensemble-based approach is in dissecting the dynamical properties of a circuit, e.g., transitions between cellular states. An archetypal system to assess RACIPE is the gene regulation of cell cycle progression because of the following reasons. First, cell cycle circuit is evolutionarily robust, as it is required by almost every cell type and organism for survival. With RACIPE, we can perturb the system to evaluate the robustness of a cell cycle GRC. Second, cell cycle dynamics involves transitions across a series of well-studied cell cycle states/checkpoints, which allows us to test whether RACIPE can capture these non-trivial dynamics using only random models. Third, there are many existing experimental evidences and data in the studies of cell cycle, which will help to validate the computational models. Fourth, a fundamental understanding of the regulatory mechanism of cell cycle will facilitate researches in areas such as cell differentiation during normal development^19–24^ and tumorigenesis in cancer^25–27^. Fifth, though there are many existing computational models for studying cell cycle networks, different models usually emphasize different aspects of the system. For example, when studying a specific checkpoint, Tyson *et al.* proposed a two-state switching model^28,29^; when studying a full cycle of the progression, multiple models considered its dynamics as oscillations^9,30^; yet, another popular model suggested the cell cycle progression as a downhill process toward the G1 global attractor11. A RACIPE analysis could provide a holistic view of the dynamical properties of a cell cycle circuit.

In this study, we analyzed a cell cycle GRC of budding yeast with RACIPE, from which we elucidated the gene regulation during cell cycle progression and the associated state transitions. To achieve this, we first generalized RACIPE for GRCs with multiple regulatory types, including transcription, signaling (such as phosphorylation), and degradation. To carefully examine oscillatory dynamics, which were considered as an essential component of cell cycle dynamics^10,31,32^, we developed a numerical algorithm to systematically detect and characterize limit cycles for a large number of models. The generalized RACIPE allowed us to examine the sequential change of circuit dynamics in cell cycle progression. We focused on discovering the features of network dynamics unbiasedly from this statistical analysis of random models. We found that the RACIPE models of the GRC allow both stable steady states, which form clusters associated with cell cycle phases, and oscillatory states, which direct irreversible state transitions along the cell cycle progression. The model predictions are largely consistent with gene perturbation data in the literature and recent single cell RNA-seq data. Furthermore, we explored possible mechanisms to understand the irreversible state transitions directly from the steady states of random models. We hope RACIPE will facilitate systems biology network modelling and generate unbiased predictions for complex systems.

## Results

There are several existing gene regulatory circuit models for the budding yeast cell cycle in the literature^11,12,30,33,34^. Based on these data, we constructed a gene regulatory circuit of 15 genes and 38 regulatory links (**Fig. 1a)**. The majority of the components of the circuit were reported in previous studies^11,35^. In addition, we made a few modifications to the circuit, as explained in the following. First, instead of including a node representing cell size, we modeled Cln3 as the input node and the effect of cell size by the production rate of Cln3. Further, we added Whi5 and its interactions, as the role of Whi5 was discovered^35^ after the publication of the original circuit model. Lastly, we added the inhibition of Cdc20 by Cdh1 as supported by multiple evidences^13,36–38^. The final circuit contains transcription factors (SBF, Whi5, MBF, Mcm1, SFF, and Swi5), major cyclin genes (Cln3, Cln1,2, Clb5,6, and Clb1,2), and the other regulatory signaling proteins (Sic1, Cdh1, DNAS, Cdc20, Pds1, and Cdc14). These genes regulate one another by either transcription (black lines), degradation (red lines), or signaling (green lines) (**Fig. 1a)**. Each interaction can be either excitatory (arrow head) or inhibitory (circle head).

**Figure 1.**
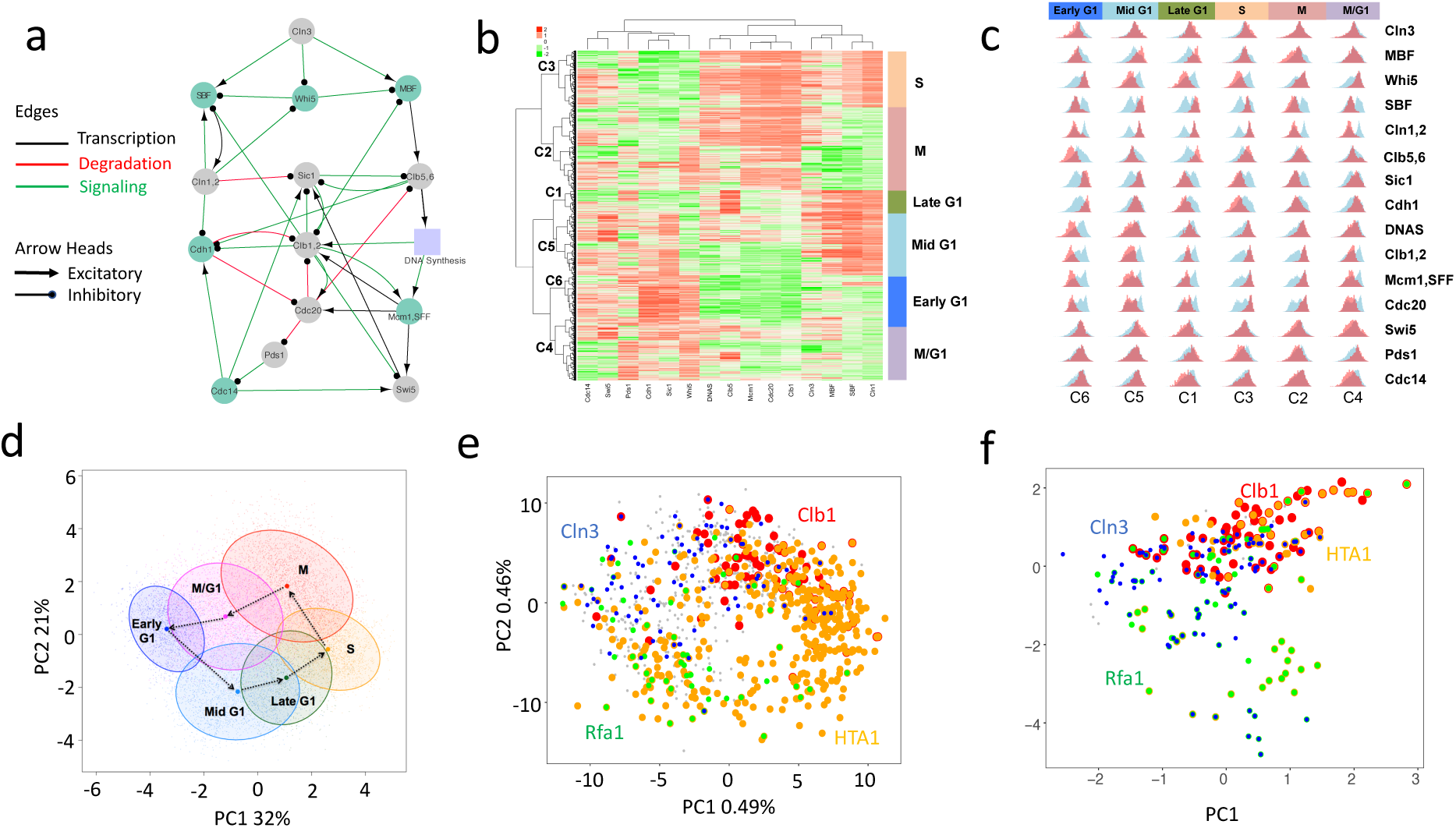
Stable steady states of the gene regulatory circuit capture cell cycle phases. (a) The core gene regulatory circuit of the yeast cell cycle. The nodes correspond to 14 genes and an abstract node representing DNA synthesis process, and the edges correspond to 38 regulatory interactions. These interactions can be any of the following three types: transcription (black), degradation (red), and signaling (green), where each type can be either excitatory (line and arrow head) or inhibitory (line and circle head). (b) Hierarchical clustering analysis were performed using 12,758 activity profiles from simulated stable steady states. Six clusters of simulated stable steady states are mapped to specific cell cycle phases. (c) The distribution of the simulated gene activity profiles of each cell cycle phase (red) and that of all the models (blue). (d) Projection of the simulated activity profiles onto the first two principal components (PCs, PC1: ∼32%, PC2: ∼21%). Each projected cluster that is associated with a cell cycle phase is presented by an ellipse whose center is illustrated by a dot. (e) Projection of the single cell RNA-seq data onto the first two PCs (PC1: 0.49%, PC2: 0.46%) obtained from the sequencing data. Cln3 expression peaks at G1 phase (blue), Rfa1 peaks (green) across late G1 and S phases, HTA1 peaks (orange) at S, and Clb1 peaks (red) at M. (f) Projection of the single cell expression data of 15 representive genes (after knn smoothing, see SI for details) onto the first two PCs from the simulated data (same as d).

To elucidate the dynamics of gene regulation during cell cycle progression, we applied a generalized version of our mathematical modeling algorithm RACIPE to the cell cycle gene circuit. In a previous study, the Boolean network model was applied to a similar circuit^11^. In comparison, our approach (1) models the activity of each gene as a continuous variable; (2) considers the differences in various regulatory types; (3) systematically characterizes both stable steady states and oscillatory dynamics (see Methods for details). We generated 10,000 RACIPE models, from which we found 12,758 stable steady states (∼84% models), and 3,070 stable oscillatory states (∼24% models). About 9% models allow the coexistence of both dynamical modes and another 1% are nonconvergent.

### The stable steady states of the circuit capture cell cycle phases

We first investigated the stable steady states from the RACIPE models. Here, the simulated activity levels of 15 genes from the 12,758 states were first log-transformed and standardized. The data is then analyzed by unsupervised hierarchical clustering analysis (R function *hclust* with distance metric as *Euclidean* and method *ward.D2*), from which we found six robust clusters, labeled as C1 ∼ C6 (**Fig. 1b**). To characterize the biological meaning of these clusters, we compared the expression patterns from the simulations with the literature data as follows. First, we computed, for each gene, the distributions of the simulated activity levels for each cluster (**Fig. 1c,** red histograms) and those of all the data (**Fig. 1c,** blue histograms). Second, we assigned the states of each gene to either high, medium, or low by comparing the red and blue histograms (**Supplementary Table 3**). Third, we compared the expression pattern of each cluster with the experimental evidence in the literature and associated each cluster to the cell cycle phase that has the best matching (**Supplementary Table 4**). The assignment of cell cycle phases for early G1, mid G1, late G1, S, M, and M/G1 is illustrated in **Fig. 1b, c.**

Next, we projected the simulated steady states to the first two principal components (∼32% contribution from PC1 and ∼21% from PC2). As shown in **Fig. 1d**, the six clusters of the steady states are arranged circularly. Strikingly, the directionality of cell cycle progression follows a series of consecutive state transitions from a cluster to the neighboring cluster along the counterclockwise direction. We then compared the simulation results with recently published single cell RNA-seq data of budding yeast39. When principal component analysis (PCA) was performed on the genome-wide gene expression data, we also observed a similar circular structure of gene expression for individual cells (Fig. 1e). Representative marker genes Cln3 (Early G1), Rfa1 (Late G1), HTA1 (S), and Clb1 (M) have peak expression in the corresponding cell cycle phases. Subsequently, we projected the expression data of genes associated with the cell cycle GRC to the PCs obtained from the RACIPE simulations. We found one of the genes (Whi5) in the GRC is not expressed at all and another gene (Cdc14) is not highly expressed in the phases it is supposed to be expressed. In those cases, we replaced the gene with another gene that (1) directly interacts with the original gene and (2) has expression consistent with previously reported microarray data^40^. Because of the dropout effect, we also performed k-nearest neighbors smoothing41 to the scRNA-seq data. As shown in **Fig. 1f**, although much noisier, the circular structure can also be discerned even with the projection of only fifteen genes (detailed procedure in Supplementary Note 1). Our finding that cell cycle progression corresponds to the state transitions in a circular structure has also been observed in budding yeast time series microarray data^40,42^ and mouse embryonic stem cells single cell RNA-seq data^43^.

The aforementioned mapping of the cell cycle phases helps to identify, from RACIPE models, the dynamical change of the gene activities along cell cycle progression. As shown in **Fig. 1c**, the histogram of the activity of some genes (e.g., SBF) is bimodal, and the gene switches the level of its activity discretely; while some other genes (e.g., Cdh1) have more continuous changes in the activity during state transitions. Markedly, the three cyclin genes Cln1,2, Clb5,6, and Clb1,2 belong to the former type. Our model predicts that Cln1,2 switches from the low to high activity in the mid G1 phase remains high till the S phase, and then switches back to low activity in the M phase. By constrast Clb5,6 switch to high in the late G1 phase, while Clb1,2 switch to high in the S phase. The model predictions on the discrete changes of the cyclin expressions and their specifc order are consistent with known experimental observations^40,44^.

### Perturbation analysis identifies the role of the genes in cell cycle progression

To further validate the modeling results of the cell cycle circuit, we performed *in silico* perturbation analysis to evaluate the effects of gene knockdown (KD) on the behavior of the circuit. The outcomes of the gene perturbations were then compared with experimental evidence in the literature. **Fig. 2** shows the modeling results for the untreated (UT), 15 single gene KD, and 12 double gene KD. Here, the effect of gene KD was simulated by reducing the maximum production rate of the corresponding genes by 95%. For each case, we simulated 10,000 random models by RACIPE and evaluated the distribution of the stable steady states in each cell cycle phase. To rapidly assign the cell cycle phase to every stable steady state, we trained a fast forward neural network model (see Methods for details) using the original 10,000 RACIPE models (i.e., the UT condition, the data from **Fig. 1b**). From our benchmark, the neural network model has over 90% accuracy in the state assignment (**Supplementary Fig. 14**).

**Figure 2.**
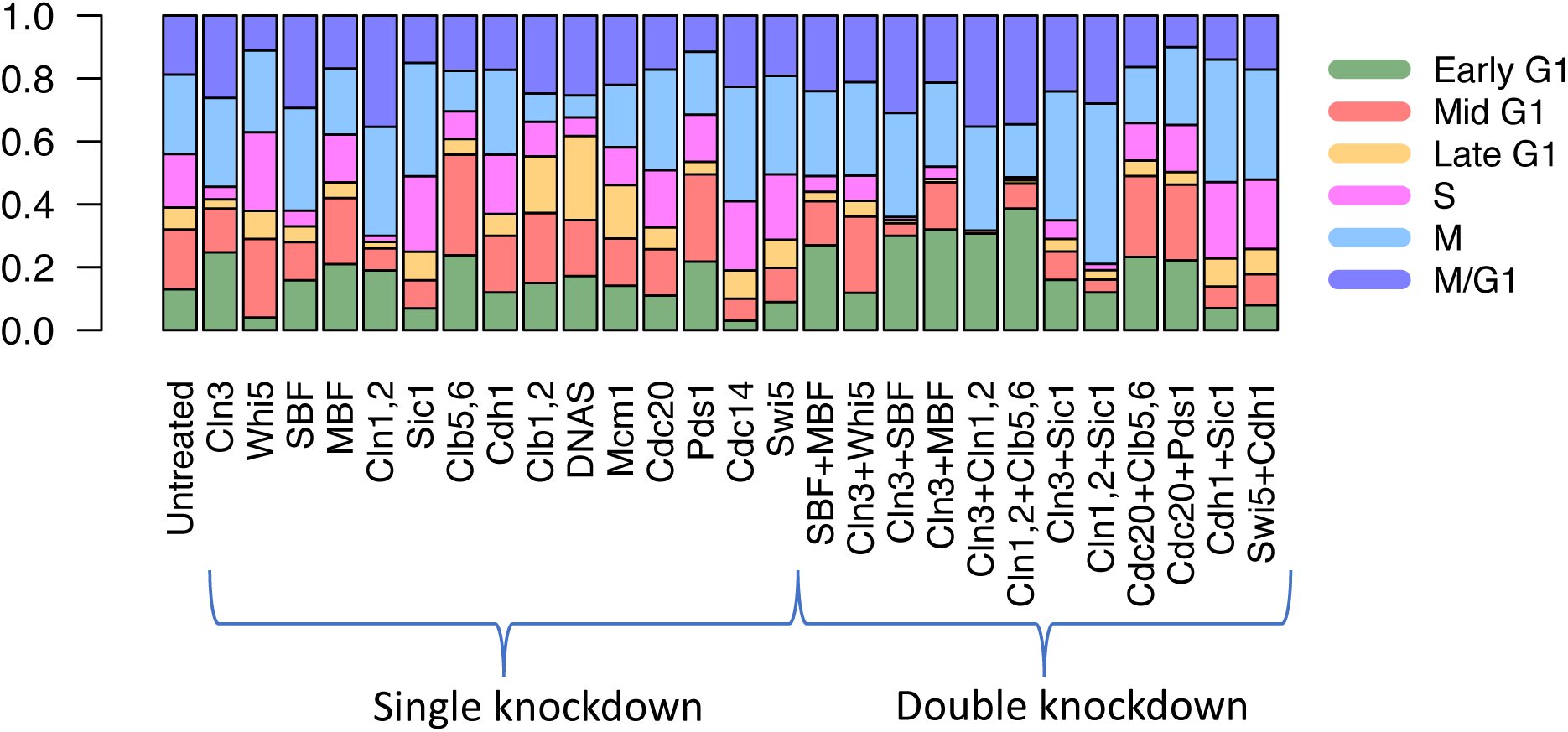
Perturbation analysis. Distribution of the fraction of steady states in each cell cycle phase from the simulations of the cell cycle circuit under untreated (UT), single knockdown (KD), and double KD conditions. To assign new phases after a perturbation, we trained a neural net model using the activity profiles of the untreated condition and the classification of cell cycle phases from the previous hierarchical clustering analysis (see Methods for details), and then we infer the phase for any new activity profile using the trained model.

The perturbation analysis identifies the role of each gene in stabilizing or destabilizing various cell cycle phases. The KD conditions where one or multiple cell cycle phases are mainly affected are summarized in **Supplementary Fig. 4 and Supplementary Table 5.** The outcomes of the gene perturbation are well supported by experimental evidence. For example, (1) our modeling results show that Cln3 KD increases the fraction of states in early G1 and M/G1 and decreases that of S. This is consistent with the experimental data where the inactivation of Cln3 was found to cause the cell size to increase, elongating early G1^45–48^. For the cell cycle START to take place, activation of Cdc28 by G1-cyclins Cln1, Cln2, or Cln3 is necessary^45^, and Cdc28 mutants cause a cell to stall at the START of early G1 phase^49^. (2) Whi5 KD increases the fraction of states in mid G1 and S and decreases that in M/G1 and early G1, consistent with the finding that the inactivation of Whi5 dramatically reduces the cell size by shortening early G1^50^. (3) Interestingly, Whi5 KD seems to have the opposite effect of Cln3 KD. Double perturbation of Cln3 and Whi5 appears to mitigate the impact of each other by restoring the proportions in early G1 and mid G1 which is consistent with experimental findings^50^. (4) Cln1 and Clb5 double KD reduces the proportion of stable states in mid G1 to M and increases that in M/G1 and early G1. This prediction is also supported by the experimental finding that quadruple knockdown of Cln1,2 and Clb5,6 arrest cells in G1^51^. (5) Cdc20 and Pds1 double KD decreases the proportion of states in M/G1, consistent with the findings that double mutant (Cdc20 and Pds1) cells get arrested in telophase^52^. These experimental evidences are listed in tabular form in **Supplementary Table 6.**

### Dynamical changes of circuit states along cell cycle progression

As shown above, stable steady states from RACIPE models can capture the gene activity patterns of various cell cycle phases. Next, we show, from these steady states altogether, how we can recapitulate the dynamic behavior of the cell cycle circuit. Based on the results of hierarchical clustering analysis in **Fig. 1b**, we assigned pseudotime index to each activity profile of a steady state (see **Supplementary Note 3** for details). The change in gene activity for Whi5 and three cyclins (Cln1,2 – the G1 phase cyclin, Clb5,6 – the S phase cyclin, and Clb1,2 – the M phase cyclin) along the pseudotime are shown in **Fig. 3a**. The dynamic change of the other genes is shown in **Supplementary Fig. S2a**. The profiles are illustrated with two repeats to represent two cycles, and they capture the overall gene activity dynamics well. Interestingly, genes like Cln1,2 exhibit binary activity – Cln1,2 level is high during G1 and low during M and M/G1. However, genes like Whi5 and Clb1,2 exhibit more gradual changes in activity along cell cycle progression. Moreover, Whi5 and Cln1,2 have opposite phases, with Whi5 switches activity sightly ahead of Cln1,2, which is captured by their negative cross-correlation (**Supplementary Fig. S2c**). Noticeably, Clb5,6 level increases when the Cln1,2 level is still high. When Clb5,6 level decreases, Clb1,2 level starts to rise and continues to stay elevated during M/G1, as its coordination with the other genes (specifically Sic1, Cdh1, Cdc14, and Swi5) is needed for mitotic exit^53–55^. The synchrony of Sic1 and Cdh1 with Clb1,2 is further observed in their autocorrelations (**Supplementary Fig. S2b**), emphasizing the role of Clb1,2 in mitotic exit.

**Figure 3.**
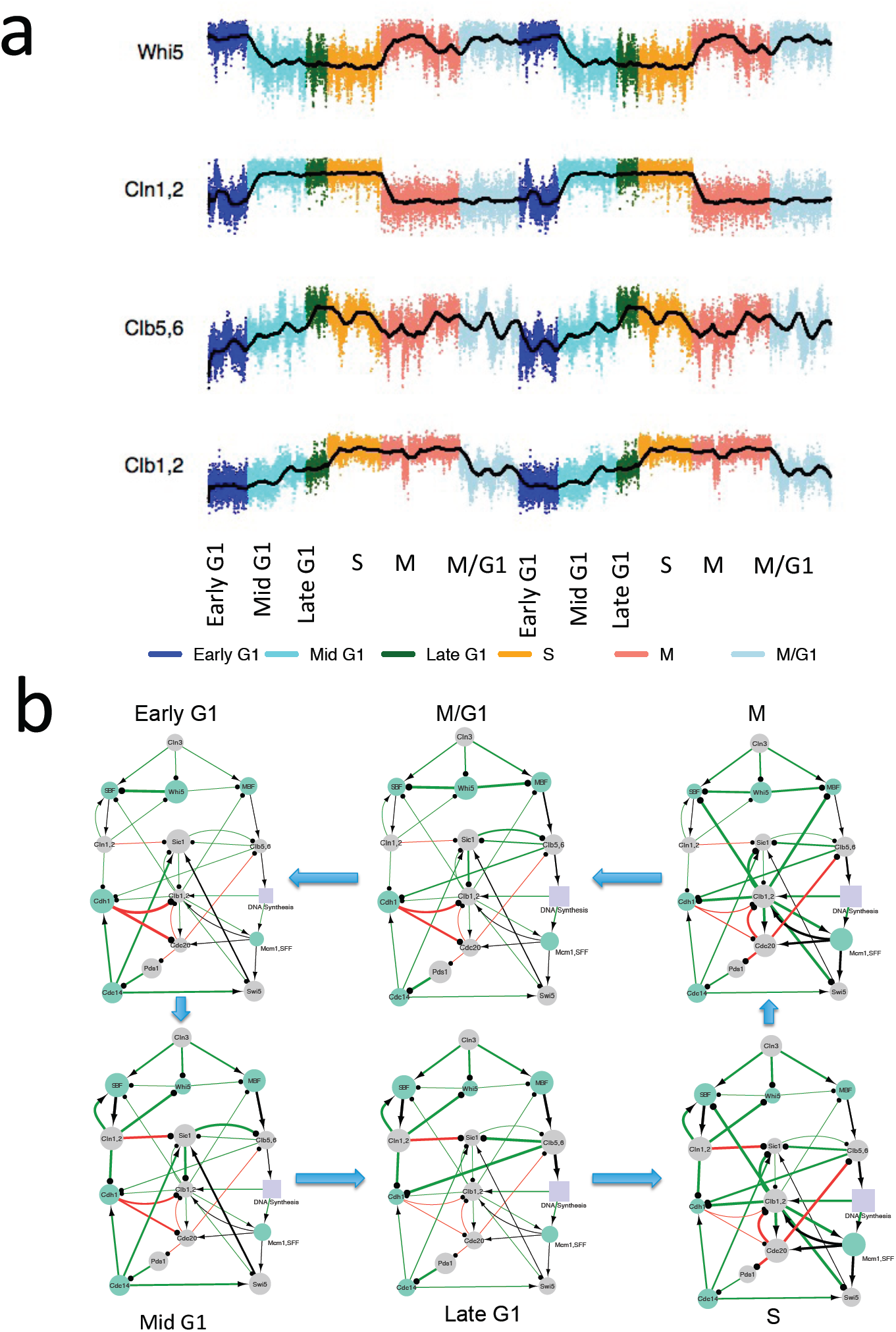
Dynamic change of gene expressions during cell cycle progression. (a) The activity levels of Whi5, Cln1,2, Clb5,6, and Clb1,2 along the inferred pseudotime (with two cycles). Colors annoate different cell cycle phases. (b) A dynamic network view of cell cycle progression. The size of each node represents the level of gene activity, and the thickness of the each edge represent the activity of the regulation, defined as the proportion of models where the level of the regulator is higher than the threshold level of the edge.

Coordinated activation of a subset of genes and their interactions drive the cell cycle from one phase to the next along the cell cycle progression. Using the simulated data from the stable steady states of 10,000 RACIPE models, we can visualize the dynamic changes of the cell cycle gene circuit (**Fig. 3b**). For each cell cycle phase, the size of a node is scaled to the average activity level of the corresponding gene from all the models in that phase; while the thickness of an edge is scaled to the fraction of models where the activity of the regulator is larger than the threshold of the Hill function (see Methods for the mathematical equations). Remarkably, this representation provides a dynamic view of gene regulation along cell cycle progression. In early G1, Whi5 has high activity which inhibits SBF, and consequently keeps Cln1,2 activity suppressed. An increased level of Cln3 suppresses Whi5 and enables SBF release. The increased level of SBF activates Cln1,2, which induces the transition to mid G1. Further, high level of Cln1,2 inhibits Whi5 and Sic1, which transitions the cell to enter late G1. This positive feedback of Cln1,2 behaves like a switch. This feedback mechanism is known as the START checkpoint, whose activation makes the cells commit to the S phase (DNA synthesis)^56,57^. In contrast to models where one network is associated with one state, here we show that a dynamic network model can describe the sequential changes of the gene regulation in all the cell cycle phases.

### Oscillatory dynamics specify the directionality of the cell cycle state transitions

In the previous sections, we analyzed the steady states of RACIPE models and associated them with various cell cycle phases. However, the analysis so far doesn’t explain the direction of transitions between these phases. Here, we explore how the state transitions can be understood from the oscillatory states of RACIPE models.

To facilitate the statistical analysis on oscilations, we obtained 47,637 oscillatory states from 160,000 RACIPE models (10,000 models from the prior analysis plus 150,000 models generated anew). For each oscillation, we projected the trajectory to the first two principal components of the steady state data (same PCs as in **Fig. 1d**). Some of the projected trajectories reside in one or a very few cell cycle phases, while some others can span many phases (**Fig. 4a**). To characterize the transitions through various cell cycle phases along an oscillatory trajectory, we used the previously trained neural network model to predict the cell cycle phase for any given gene activity profile along an oscillatory trajectory. This allows us to identify the patterns of state transitions for each oscillation. For example, two oscillations are shown in **Fig. 4a**: the one on the left shuffles back and forth between two phases (M → M/G1 → M), whereas the one on the right passes through all the phases (early G1 → mid G1 → late G1 → S → M → M/G1 → early G1).

**Figure 4.**
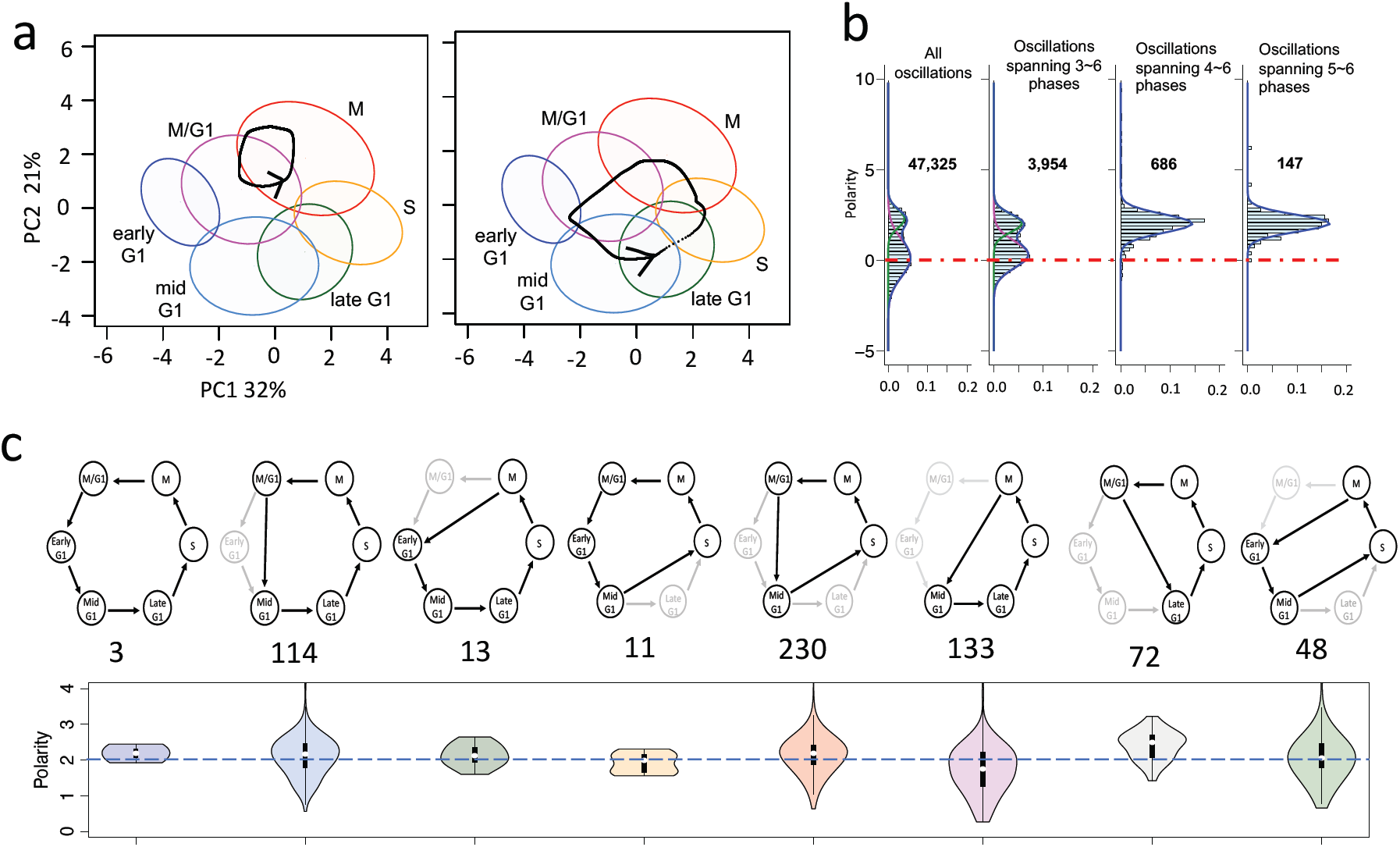
Statistical analysis of oscillatory trajectories elucidates cell cycle dynamics. **(a)** Examples of the oscillatory trajectories projected on the first two principal components (PC1 and PC2) obtained from the stable states from 10,000 models. Left panel: Trajectory travelling only two cell cycle phases; right panel: Trajectory traveling all six cell cycle phases. An arrow on the projected trajecgtory marks the direction of oscillation. **(b)** The histogram of the polarity of oscillatory trajectories of different sizes – all trajectories (n = 47,327), trajectories spanning 3∼6 phases (n = 3,954), 4∼6 phases (n = 686), and 5∼6 phases (n = 147). The histograms are fitted by either two Gaussian distributions (first two columns, where green curve for the distribution with a positive mean, pink curve for the distribution with a near zero mean) or a single Gaussian distribution (the last two columns). The horizontal dotted red line shows the zero polarity. **(c)** Top panel shows eight major state transition patterns of oscillatory trajectories. The number below each pattern indicates the number of observed oscillations. Bottom panel shows the histogram of polarity of the each pattern. Data for the other recurrent patterns are shown in Supplementary Fig. S3

To quantify the directions of the oscillatory trajectories, we defined an angular-velocity-based metric, named as polarity, to measure whether a projected oscillatory trajectory is clockwise or counterclockwise (see Methods). A positive polarity corresponds to a counterclockwise trajectory. Interestingly, the distribution of the polarity for all the projected trajectories is bimodal (**Fig. 4b**, leftmost panel), which can be fit nicely by two Gaussian distributions – one with the mean zero (pink curve) and the other with positive values for almost the whole distribution (green curve). However, if we considered the trajectories spanning four or more cell cycle phases, the distribution of the polarity can be fit by a single Gaussian distribution with only positive values (**Fig. 4b**, two rightmost panels). This result suggests that the polarity for the trajectories of small size (i.e., spanning no more than three phases) is random, presumably because there is no restraint for the oscillations to travel. But the polarity for the trajectories of large size (i.e., spanning four or more phases) is mostly positive. These oscillations only travel counterclockwise, consistent with the known sequence of the cell cycle progression. Note that we found this consistent pattern of state transitions from randomly generated models because the gene circuit topology restrains the possible routes of state transitions. **Fig. 4c** lists eight major recurring transition patterns involving four or more phases with positive polarity i.e. all the corresponding oscilations travel counterclockwisely. The plolarity distributions of the other transition patterns are shown in **Supplementary Fig. S3**.

### Cell cycle gene circuit specifies irreversible phase transitions

So far, we have shown that the cell cycle phases can be described by the stable steady states from RACIPE models, while the directionality of the cell cycle progression can be deduced from the oscillatory states. However, there is a remaining issue in this theory of directionality of cell cycle progression as follows. RACIPE models with random kinetic parameters can be regarded as cells under different signaling or environmental conditions. Thus, it makes sense that different models can correspond to different cell cycle phases. However, if the signaling conditions can be altered reversibly, the state transitions should also be reversible, which is not consistent with the fact that the cell cycle progression is irreversible. To address this issue, we propose the following two analyses to explain the irreversible state transitions in the cell cycle using the stable steady state solutions from the RACIPE models.

First, we show in the following that the irreversible state transitions can be inferred using the steady state gene activity profiles and the topology of the cell cycle gene circuit. The approach is based on the fact that the activity of the targets lags behind the activity of the regulators. Thus, for a given transition from state 1 to state 2, the activity correlation of the regulators in state 1 and the targets in state 2 should be higher than the activity correlation of the regulators in state 2 and the targets in state 1. Therefore, we evaluate a so-called *delayed correlation* for the forward and backward transitions between state 1 and state 2 (see Methods for details). If the median delayed correlation is significantly different between the forward and backward directions, the state transition is likely to be irreversible. As shown in **Fig. 5a,b** and **Supplementary Fig. S4**, the delayed correlations for the forward transitions of consecutive cell cycle phases are usually higher than those for the backward transitions, except for the M to M/G1 and M/G1 to early G1 transitions where both forward and backward correlations are comparable. Based on these delayed correlations, we derive a state transition model for cell cycle progression (**Fig. 5c**). Note that this approach is generally applicable to infer the paths of state transitions for a gene regulatory circuit, examples of which are shown in **Fig. 6b** for a repressilator circuit (details in **Supplementary Fig. S13)** and an induced toggle switch circuit (details in **Supplementary Fig. S16).**

**Figure 5.**
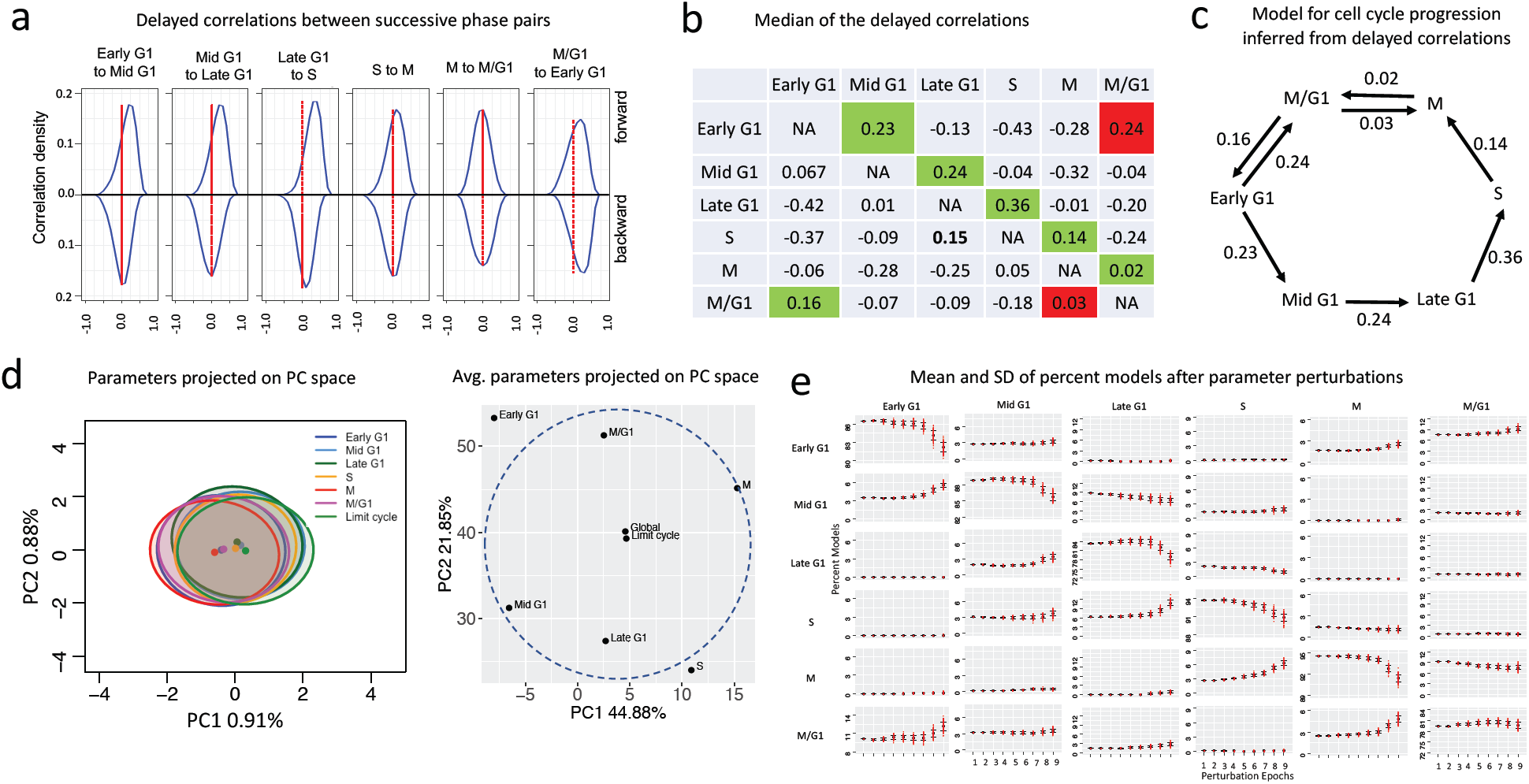
The irreversible direction of the cell cycle inferred from stable steady states. **(a-c)** A model for cell cycle progression inferred from the delayed correlations: **(a)** Densities of the forward (top) and backward (bottom) delayed correlations of gene expressions between successive cell cycle phases. Densities for all possible pairs are shown in Supplementary Fig. 4. The vertical dotted red line along zero correlation helps identify the skewness of the distributions. **(b)** Median delayed correlations for every two cell cycle phases. Medians along the direction of the cell cycle are predominantly larger (green), except for M to M/G1 and M/G1 to early G1 where the backward correlations are also comparable (red). **(c)** A model for cell cycle progression derived from the delayed correlations. **(d-e)** A scheme for cell cycle progression inferred from parameter perturbation. **(d)** Left panel shows the parameter vectors of every one-state model in each cell cycle phase and oscillatory state projected on the first two principal components in the parameter space. Right panel shows the averages of the parameter vectors of all the one-state models in each cell cycle phase and oscillatory state. **(e)** Mean and standard deviation (SD) of the percent models in each cell cycle phase when simulated with the parameters perturbed toward the global parameter average. Perturbed models from one cell cycle phase diffuse to the next cell cycle phase with one exception early G1. In early G1, the perturbed models diffuse to both mid G1 and M/G1.

**Figure 6.**
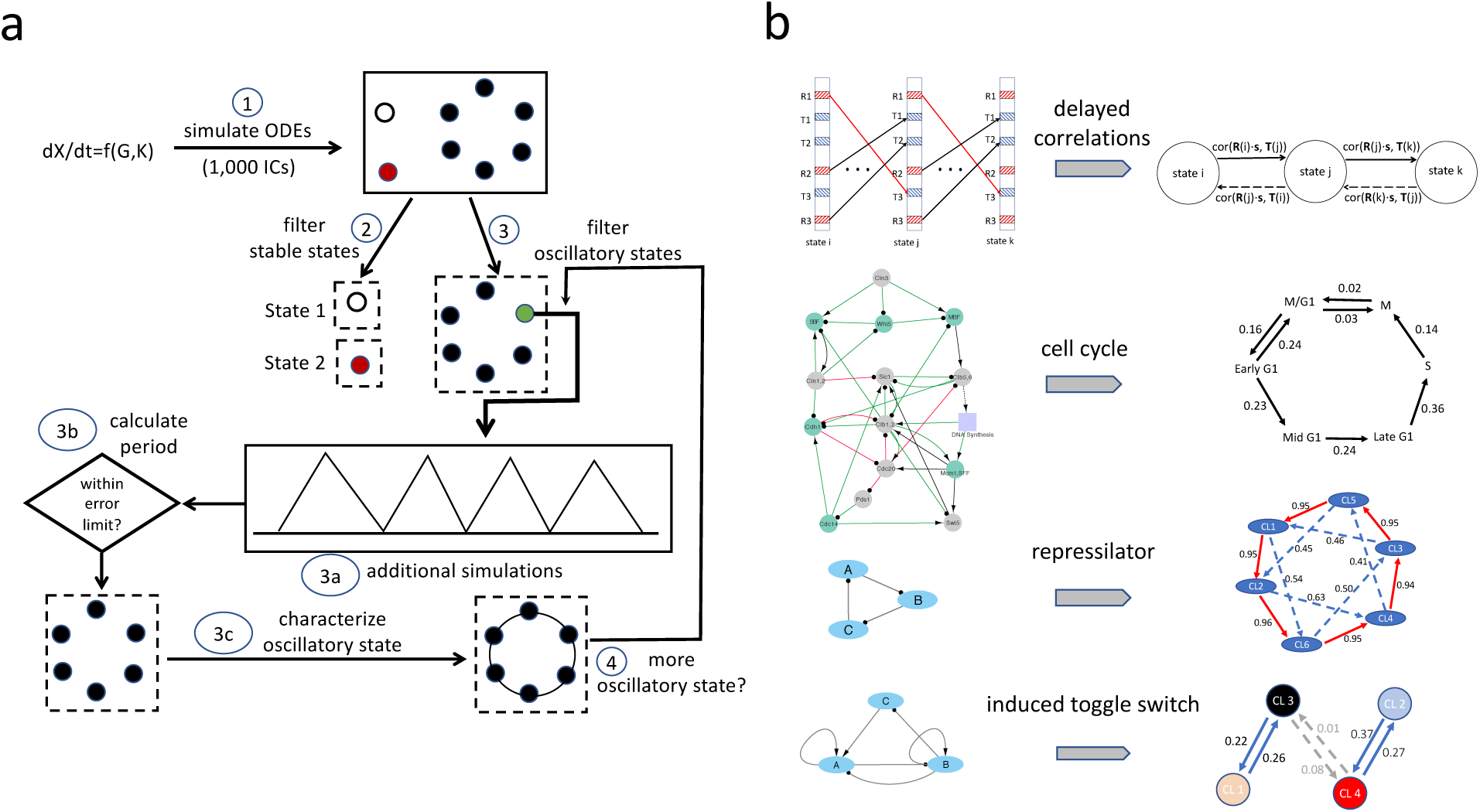
Schematic summary of the computational methods. **(a)** Flowchart for detecting the coexistence of steady states and oscillatory states in the generalized RACIPE (see Methods for details). **(b)** A method to predict state transitions using the steady state gene activity profiles of RACIPE models and the circuit topology. **1**^**st**^ **row:** schematic of the method. Left panel shows three expression/activity vectors of three states i, j, and k, respectively. A black line with an arrow head indicates an excitatory link, whereas a red line with a circle head indicates an inhibitory link, and **s**(links) is the sign vector expressing each interaction type (+1: excitatory, -1: inhibitory). Right panel illustrates the definition of the delayed correlation for forward transitions (thick black line pointed right) and backward transitions (black dotted line pointed left). 2^nd^ to 4^th^ rows show three applications of the delayed correlations to infer state transitions. **2**^**nd**^ **row:** the cell cycle circuit (left) and delayed correlation based transitions between six cell cycle phases (right). **3**^**rd**^ **row:** a repressilator circuit (left) and predicted transitions between six clusters (CL1 ∼ CL6) obtained the RACIPE models. Details are in **Supplementary Fig. S13. 4**^**th**^ **row**: a three-node induced toggle switch circuit (left) and predicted transitions between four clusters (CL1 ∼ CL4). Details in **Supplementary Fig. S16**.

Second, we propose a putative model to explain how irreversible state transitions occur by changing the kinetic parameters. In the analysis, we consider seven groups of models – each of the first six groups corresponds to the models that allow only one stable steady state of the corresponding cell cycle phase, and the seventh group corresponds to the models that generate oscillatory state spanning 3 or more cell cycle phases. We first performed PCA on the high-dimensional kinetic parameter profiles from the selected RACIPE models (# models - early G1: 635, mid G1: 689, late G1: 365, S: 721, M: 1304, M/G1: 805, and limit cycles: 3899). Unlike the gene activity profiles, the parameters from the different cell cycle phases and the oscillatory states overlap, and the magnitude of the parameters is not significantly different from each other (**Fig. 5d, left panel**). However, if PCA is performed on the average parameter profiles of each group, the projected parameter profiles follow a similar spatial arrangement as the projected gene activity profiles (**Fig. 5d right panel).** Interestingly, the average projected parameter profile for the oscillatory states is near the center and at the same time, close to the average projected parameter profile of all the models. Next, starting from the models of a cell cycle phase, we perturbed the kinetic parameters and evaluated the changes in the model proportions across the cell cycle phases. We found that, if the parameters are perturbed toward the global average, the models shift toward the next cell cycle phase for most cases, except for early G1 (**Fig. 5e, Supplementary Fig. S9**), in which models are increased in both mid G1 and M/G1. Similar observations were found if we perturbed the parameters toward the average of the oscillatory states (**Supplementary Fig. S10**), or toward the parameters of any oscillatory model (**Supplementary Fig. S11**). Note that the pattern of diffusion of the models along the cell cycle direction is not observed when the kinetic parameters are perturbed randomly (**Supplementary Fig. 12**). For details of the parameter perturbation, see **Supplementary Note 4**. Our modelling results demonstrate that the irreversible state transition during the cell cycle progression can be achieved by relaxing the kinetic parameters toward either the global average, the average of the oscillatory states, or a random model of the oscillatory states.

In short, we have shown through two different approaches how irreversible state transitions can be derived from the steady states of random models. Our simulation approach also sheds light on possible mechanisms to achieve irreversible state transitions by changing the signaling states (represented here by the kinetic parameters) of the circuit.

## Discussion

In this study, we used the budding yeast cell cycle as a model system to elucidate dynamic behaviors of gene regulatory circuits using a computational systems biology approach. We generalized and applied our recently developed mathematical modeling algorithm, named *ra*ndom *ci*rcuit *pe*rturbation (RACIPE), to analyze the dynamic features of a budding yeast cell cycle gene circuit using an ensemble of chemical rate equation models with random kinetic parameters. From an extensive statistical analysis on a large set of simulated data, we found the clusters of steady state gene activities can be associated with specific cell cycle phases, while the oscillatory states specify the path of the state transitions during the cell cycle. These cell cycle specific clusters from circuit simulations are consistent with budding yeast single cell RNA-seq data. Finally, we elucidate the mechanism of the irreversible state transitions using RACIPE-generated models. Our analyses demonstrate the role of the circuit topology in determining the cell cycle dynamics and its function. The unique combination of modeling and statistical data analysis can be a powerful approach to analyzing the behavior of complex dynamical systems in biology. Note that, because of the evolutionary conservation, most of the genes and interactions in the circuit of the budding yeast cell cycle have counterparts in mammalian systems (**Supplementary Table 2**)^58^. Thus, the analysis, although focused on budding yeast, can be extended to study the mammalian cell cycle.

Cell cycle is an ideal testing ground for mathematical modeling, because (1) there are rich datasets and literature on gene regulation in cell cycle; (2) substantial efforts have been made to model the dynamics of cell cycle gene regulatory circuits; (3) cell cycle involves transitions of multiple cell cycle phases. Thus, the dynamics are non-trivial. Elucidating cell cycle gene regulation can contribute to a better understanding of the role of the cell cycle in sundry biological processes, such as cell differentiation^19–24^, DNA damage^59–62^ and metabolism^63–65^, and in diseases, such as cancer^25–27^. Compared to previous modeling studies^11,28–30^, we have generated a large number of models and performed extensive data analysis on all of these models. Our results illustrate that models with different sets of parameters can be associated with either one of the cell cycle phases and checkpoints, or the oscillatory dynamics across multiple cell cycle phases. Strikingly, we show that the cell cycle dynamics can be well explained even without carefully selected kinetic parameters, again demonstrating the crucial role of the cell cycle gene circuit, other than the detailed kinetic parameters, in determining the circuit’s function.

To study the cell cycle gene circuit, we have improved the original RACIPE method^15^ and further developed statistical methods to analyze the simulated data from a large set of models. Primarily, RACIPE is generalized to model three additional types of regulation (phosphorylation, signaling, and degradation) in addition to transcriptional regulation. Futhermore, the method is extended for high throughput analysis of not only stable steady states, but also oscillatory dynamics. To carefully characterize different dynamical features from many models, we developed numerical algorithms to automatically detect oscillatory trajectories and characterize the state transition patterns using machine learning. The advances in RACIPE enables us to identify robust dynamical features of the cell cycle directly from randomly generated models.

An important question we focused on is to understand how the cell cycle gene circuit controls the irreversible state transitions. As shown in our simulated data, we have investigated it from three different aspects. First, we systematically mapped the patterns of state transitions from all of the RACIPE-generated oscillatory trajectories. From this, we identified not only a consistent directionality of the state transitions, but also recurring transition patterns. The statistical analysis indicates the crucial role of the circuit topology, instead of a specific set of kinetic parameters, in determining the irreversible nature of the state transitions during the cell cycle. Second, we developed a numerical method to infer the directions of state transitions directly from the steady-state gene expression/activity profiles and the network topology. Here we achieved this by considering that the expression of the targets occurs later than the activation of the regulators and employed a metric based on delayed correlations. We have applied this approach on several classic gene regulatory circuits, including the cell cycle circuit, and successfully reproduced the patterns of state transitions. The success of this method demonstrates again how gene regulatory circuit drives state transitions. Third, we examined the relationship between the cell cycle progression and the signaling states of the gene regulatory circuit by perturbing the model parameters belonging to each cell cycle phase and assessing how the models diffuse to the other phases. We perturbed the parameters either toward the global parameter average (i.e., making the parameters more balanced) or relaxed them toward the limit cycle parameters. We found that the perturbed models transition to the next cell cycle phase in both cases. This finding suggests that an irreversible state transition of the cell cycle would occur by systematically changing the signaling state of the circuit.

Overall, we have shown the power of RACIPE in gene circuit modeling. Yet, there are several issues that need to be addressed in the future. First, the state assignment is currently based on the hierarchical clustering analysis of the steady state activity profiles from the random models. The quality and robustness of the clustering might affect the performance of the subsequent statistical analysis. For instance, we have noticed a few distinct features for the M/G1 phase, which might suggest a hybrid nature of the phase. From the performance of neural net model in assigning phases, we observed that, the M/G1 phase was incorrectly assigned to the early G1 and M phases, each being at 7.59% and 6.62%, respectively; while the early G1 phase was incorrectly assigned to M/G1 by 11.99%. This state assignment problem for M/G1 suggests that the gene activities of M/G1 are similar to those of both the early G1 and M phases. Besides, in our state transition map (**Fig. 5c**), we predicted two bidirectional transitions (M vs. M/G1 and M/G1 vs. early G1), both involving M/G1. Additionally, we know that the mitotic exit network gets activated during the M/G1 phase^53–55^ which makes the activity profile of the dividing cells very similar to that of early G1 (shown in **Fig. 3b**). Second, although we have tested several cell cycle circuits with minor variations, a systematic approach for circuit refinement is needed for better modeling. One potential approach is to apply RACIPE on different versions of a circuit by adding/removing genes/links and analyzing the circuits iteratively. During the process, the modeling results from different circuits can be compared against literature or genomics data (such as gene expression data). Thus, RACIPE can not only be used for modeling existing gene circuits but can also be utilized in circuit refinement.

## Methods

### Generalized RACIPE

We have recently developed a computational algorithm, named *ra*ndom *ci*rcuit *pe*rturbation (RACIPE), for modeling the dynamics of a gene regulatory circuit without the need for specific kinetic parameters15. Compared to the conventional modeling methods, RACIPE uses the topology of a circuit as the only input for modeling. Instead of fitting kinetic parameters against experimental data (such as gene expression profiles), RACIPE generates an ensemble of mathematical models with distinct random kinetic parameters. The modeling results are then derived from the statistics of the dynamic behavior of the whole ensemble. Details of the RACIPE framework are described in one of our previous papers^14^.

In the original RACIPE, we mainly focused on modeling gene circuits with transcription factors involving transcriptional regulation and analyzing the stable steady state solutions from a random model. However, the yeast cell cycle circuit contains not only transcriptional regulation but also signaling regulation by kinases and the regulation of degradation by specialized protein complexes, such as anaphase-promoting complex widely known as APC. The cell cycle is also frequently modeled by oscillatory dynamics. To adapt the RACIPE-based approach, we have generalized the original method to allow modeling circuits with the additional types of regulatory interactions and developed a numerical method to systematically detect oscillatory states.

### Modeling multiple interaction types

In the original RACIPE, transcriptional regulation of gene *Y* to gene *X* is modeled by

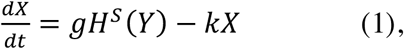

where *g* is the basal transcription rate of *X, k* is the degradation rate of *X*. *H*^*S*^ is the shifted Hill function, 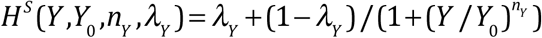, where *Y*_*0*_ is the threshold level of *Y, n*_*Y*_ is the cooperativity of the regulation from *Y* to *X*, and λ_*Y*_ is the fold change.

For a gene *X* that is regulated by signaling gene *S*, e.g., phosphorylation, we model the *activity* (instead of the *expression*) of gene *X* by

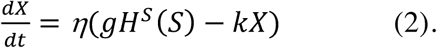

Here, we use the Hill function to model signaling regulation. As it is known that signalling transduction generates ultra-sensitivity^66^, a scaling factor *η*, an adjustable parameter included to the rate equation to account for faster time scaling of signaling regulation. For modeling the circuits used in this paper, we set this scaling factor as 10^67^. For a gene *X* whose protein degradation is regulated by gene *Z*, e.g., phosphorylation, we model the protein level of gene by

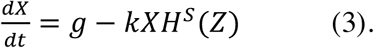

When a combinatorial interaction of multiple types is involved, e.g., transcriptional regulation by *Y*, signaling regulation by *S* and degradation regulation by *Z*, we model the activity of *X* by

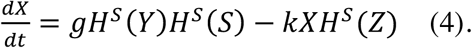

We do not include the scaling factor *η* in this situation, as transcription and/or degradation regulations are usually the rate-limiting step.

### Detecting both stable states and oscillatory states

The original RACIPE detects stable steady states of each model by first simulating ordinary differential equations (ODEs) from each of 1,000 initial conditions until the system converges to a stable steady state, and then identifies all unique steady states. It is challenging to detect oscillatory dynamics robustly and efficiently with a high-throughput numerical method, as an oscillatory system never converges to a constant expression. Additionally, we need to detect the coexistence of both the stable steady states and the oscillatory states. Here, we design and implement a fast-numerical method that automatically detects both oscillatory states and stable states (**Fig. 6a**). For each model, we first simulate the ODEs, starting from each of the 1,000 initial conditions for a fixed duration. The algorithm detects the stable-state solutions as well as other solutions that fail to converge to stable states. For the former solutions, we identify all unique stable steady states. The latter situations could arise because of either a truly non-convergence condition or an oscillation or other complex trajectory. Therefore, for the latter case, we detect whether it is an oscillation by running the simulation further and checking for recurrence at periodic intervals. If it is an oscillation, we compute its period. We repeat this process for other initial conditions and discover all the oscillatory states with distinct periods. Further details are in **Supplementary Note 2**.

### A polarity metric of limit cycles

For an oscillatory state, we project the corresponding oscillatory trajectory to the first two principal components (PCs) that are derived from all of the stable steady states from the 10,000 RACIPE models. In this two-dimensional space, we quantify the directionality of a projected oscillatory trajectory (limit cycle) by a polarity metric, defined below:

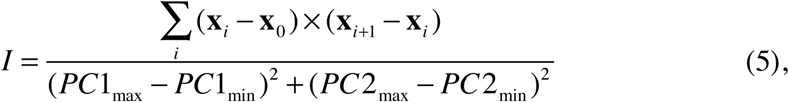

where the denominator is the sum of the squares of each extent of the limit cycle along PC1 and PC2 (PC1_min_ and PC1_max_ are the minimum and the maximum of the limit cycle along PC1, and PC2_min_ and PC2_max_ are the minimum and the maximum along PC2) and the numerator is the sum over each magnitude of the cross product between the vector pair x_i_-x_0_ and x_i+1_-x_i_ where x_0_ is the position vector for the center of the limit cycle, x_i_ is the position vector for a point i where i runs through each point along the limit cycle. The center of the limit cycle was obtained by projecting the mean expression of all the expression vectors along the oscillatory trajectory on the first two PCs. A positive polarity indicates a counterclockwise oscillation of the projected trajectory. Likewise, a negative polarity corresponds to a clockwise oscillation.

### Identifying the state transitions in an oscillation

To further characterize the directionality of an oscillatory trajectory without the dimension reduction by PCA, we use the activity profile along the oscillatory trajectory and adopt a machine learning approach on these data to identify the cell cycle phases traveled by the trajectory. From this sequence of phases, we can obtain the pattern of state transitions for each oscillation. First, we apply *nnet*^68^ to train a feed-forward neural network using the activity profile of 10,000 RACIPE models (a total of 12,758 stable steady states), where the data are classified by their cell cycle phases obtained from the hierarchical clustering analysis (**Fig. 1b**). Second, we use the trained model to assign cell cycle phases to the points along the oscillatory trajectory. Here, a data point is assigned to a cell cycle phase with the highest probability from the neural network. To achieve high accuracy in the prediction, we only make the assignment to the points when the differences between the first two highest probabilities are larger than a threshold value T. The remaining points are not included in the calculation of the cell cycle phase transition pattern of the oscillatory trajectory.

We tested the performance of the neural network model by employing a 10-fold cross-validation on the data set of 12,758 states. The performance was evaluated for data points that are retained after the filtering step. The first two panels of **Supplementary Fig. S14** show the accuracy of the model on the training and testing sets for various T threshold values. It is clear that the accuracy consistently gets better as T increases. However, the coverage (defined as the percentage of retained points) decreases as T increases. We chose T as 0.25, as it provides a right balance of accuracy and coverage, both being around 91%.

After assigning the cell cycle phases to the points along a limit cycle trajectory, we further obtain the cell cycle phase transition pattern by removing the redundant phases from every consecutive repeat (which represents more cycles). With this strategy, we compute the unique state transition patterns of all the oscillatory trajectories (47,637) from the 160,000 RACIPE models. We find 47 unique patterns from the trajectories passing through 3 or more cell cycle phases. A list of these patterns and their frequencies can be found in **Supplementary Table 7**.

### Determining directionality from circuit-topology-based delayed correlations

We further developed a correlation-based method to predict the propensity of transition between two cell cycle phases using the RACIPE-generated simulated gene activity data and the circuit topology. In this method, we calculate the delayed correlation of each pair of cell cycle phases as follows. We compute the Pearson’s correlations between the gene activities of two models where the first model coming from the first cell cycle phase 1 and the other model from the other phase 2. The forward correlation between phase 1 and phase 2 is cor(**T**(2)×**s**(links), **R**(1)), where **T**(2) is the activity vector of the targets belonging to the phase 2 model, **R**(1) is the activity vector of the sources/regulators belonging to the phase 1 model, and **s**(links) is the sign vector expressing each interaction type (+1: excitatory, -1: inhibitory). Thus, the backward correlations for the same model pair can be calculated as cor(**T**(1)×**s**(links), **R**(2)). To infer the direction of state transitions, we evaluate the median delayed correlations for any pair of models from the two phases – significantly larger (positive) forward correlation suggests an irreversible transition from phase 1 to phase 2.

## Code availablity

The software for the generalized RACIPE will be made publicly available.

## Acknowledgments

The study is supported by a startup fund from The Jackson Laboratory, by the National Cancer Institute of the National Institutes of Health under Award Number P30CA034196, and by the National Institute of General Medical Sciences of the National Institutes of Health under Award Number R35GM128717.

## REFERENCES

1. Gerstein, M. B. et al. Architecture of the human regulatory network derived from ENCODE data. Nature 489, 91–100 (2012).

2. Schwikowski, B., Uetz, P. & Fields, S. A network of protein-protein interactions in yeast. Nat. Biotechnol. 18, 1257–1261 (2000).

3. Saelens, W., Cannoodt, R., Todorov, H. & Saeys, Y. A comparison of single-cell trajectory inference methods. Nat. Biotechnol. 37, 547 (2019).

4. Moignard, V. et al. Decoding the regulatory network of early blood development from single-cell gene expression measurements. Nat. Biotechnol. 33, 269–276 (2015).

5. Farrell, J. A. et al. Single cell reconstruction of developmental trajectories during zebrafish embryogenesis. Science 360, eaar3131 (2018).

6. Ciliberti, S., Martin, O. C. & Wagner, A. Robustness can evolve gradually in complex regulatory gene networks with varying topology. PLoS Comput. Biol. 3, e15 (2007).

7. Balleza, E. et al. Critical dynamics in genetic regulatory networks: examples from four kingdoms. PLOS ONE 3, e2456 (2008).

8. Sanchez-Osorio, I., Ramos, F., Mayorga, P. & Dantan, E. Foundations for modeling the dynamics of gene regulatory networks: a multilevel-perspective review. J. Bioinform. Comput. Biol. 12, 1330003 (2013).

9. Csikász-Nagy, A., Battogtokh, D., Chen, K. C., Novák, B. & Tyson, J. J. Analysis of a Generic Model of Eukaryotic Cell-Cycle Regulation. Biophys. J. 90, 4361–4379 (2006).

10. Goldbeter, A. et al. From simple to complex oscillatory behavior in metabolic and genetic control networks. Chaos Interdiscip. J. Nonlinear Sci. 11, 247–260 (2001).

11. Li, F., Long, T., Lu, Y., Ouyang, Q. & Tang, C. The yeast cell-cycle network is robustly designed. Proc. Natl. Acad. Sci. U. S. A. 101, 4781–4786 (2004).

12. Li, C. & Wang, J. Landscape and flux reveal a new global view and physical quantification of mammalian cell cycle. Proc. Natl. Acad. Sci. 111, 14130–14135 (2014).

13. Hong, C. et al. A checkpoints capturing timing-robust Boolean model of the budding yeast cell cycle regulatory network. BMC Syst. Biol. 6, 129 (2012).

14. Huang, B. et al. RACIPE: a computational tool for modeling gene regulatory circuits using randomization. BMC Syst. Biol. 12, 74 (2018).

15. Huang, B. et al. Interrogating the topological robustness of gene regulatory circuits by randomization. PLOS Comput. Biol. 13, e1005456 (2017).

16. Kohar, V. & Lu, M. Role of noise and parametric variation in the dynamics of gene regulatory circuits. Npj Syst. Biol. Appl. 4, 40 (2018).

17. Jia, D. et al. Elucidating cancer metabolic plasticity by coupling gene regulation with metabolic pathways. Proc. Natl. Acad. Sci. 116, 3909–3918 (2019).

18. Bocci, F. et al. NRF2 activates a partial epithelial-mesenchymal transition and is maximally present in a hybrid epithelial/mesenchymal phenotype. Integr. Biol. 11, 251–263 (2019).

19. Li, V. C. & Kirschner, M. W. Molecular ties between the cell cycle and differentiation in embryonic stem cells. Proc. Natl. Acad. Sci. 111, 9503–9508 (2014).

20. Dalton, S. & Coverdell, P. D. Linking the cell cycle to cell fate decisions. Trends Cell Biol. 25, 592–600 (2015).

21. Ruijtenberg, S. & van den Heuvel, S. Coordinating cell proliferation and differentiation: antagonism between cell cycle regulators and cell type-specific gene expression. Cell Cycle 15, 196–212 (2016).

22. Soufi, A. & Dalton, S. Cycling through developmental decisions: how cell cycle dynamics control pluripotency, differentiation and reprogramming. Development 143, 4301–4311 (2016).

23. Jakoby, M. & Schnittger, A. Cell cycle and differentiation. Curr. Opin. Plant Biol. 7, 661–669 (2004).

24. Myster, D. L. & Duronio, R. J. Cell cycle: to differentiate or not to differentiate? Curr. Biol. 10, R302–R304 (2000).

25. Icard, P., Fournel, L., Wu, Z., Alifano, M. & Lincet, H. Interconnection between Metabolism and Cell Cycle in Cancer. Trends Biochem. Sci. 44, 490–501 (2019).

26. Collins, K., Jacks, T. & Pavletich, N. P. The cell cycle and cancer. Proc. Natl. Acad. Sci. 94, 2776–2778 (1997).

27. Sandal, T. Molecular Aspects of the Mammalian Cell Cycle and Cancer. The Oncologist 7, 73–81 (2002).

28. Barik, D., Baumann, W. T., Paul, M. R., Novak, B. & Tyson, J. J. A model of yeast cell-cycle regulation based on multisite phosphorylation. Mol. Syst. Biol. 6, 405 (2010).

29. Tyson, J. J. & Novak, B. Control of cell growth, division and death: information processing in living cells. Interface Focus 4, 20130070 (2014).

30. Gerard, C. & Goldbeter, A. Temporal self-organization of the cyclin/Cdk network driving the mammalian cell cycle. (2009).

31. Novák, B. & Tyson, J. J. Design principles of biochemical oscillators. Nat. Rev. Mol. Cell Biol. 9, 981–991 (2008).

32. Tyson, J. J., Chen, K. C. & Novak, B. Sniffers, buzzers, toggles and blinkers: dynamics of regulatory and signaling pathways in the cell. Curr. Opin. Cell Biol. 15, 221–231 (2003).

33. Battogtokh, D. & Tyson, J. J. A Bistable Switch Mechanism for Stem Cell Domain Nucleation in the Shoot Apical Meristem. Front. Plant Sci. 7, (2016).

34. Chen, K. C. et al. Integrative Analysis of Cell Cycle Control in Budding Yeast. Mol. Biol. Cell 15, 3841–3862 (2004).

35. Barik, D., Ball, D. A., Peccoud, J. & Tyson, J. J. A stochastic model of the yeast cell cycle reveals roles for feedback regulation in limiting cellular variability. PLOS Comput. Biol. 12, e1005230 (2016).

36. Mangla, K., Dill, D. L. & Horowitz, M. A. Timing robustness in the budding and fission yeast cell cycles. PLoS One 5, e8906 (2010).

37. Okabe, Y. & Sasai, M. Stable Stochastic Dynamics in Yeast Cell Cycle. Biophys. J. 93, 3451–3459 (2007).

38. Rudner, A. D. & Murray, A. W. Phosphorylation by Cdc28 Activates the Cdc20-Dependent Activity of the Anaphase-Promoting Complex. J. Cell Biol. 149, 1377–1390 (2000).

39. Jackson, C. A., Castro, D. M., Saldi, G.-A., Bonneau, R. & Gresham, D. Gene regulatory network reconstruction using single-cell RNA sequencing of barcoded genotypes in diverse environments. bioRxiv (2019). doi:10.1101/581678

40. Spellman, P. T. et al. Comprehensive identification of cell cycle–regulated genes of the yeast Saccharomyces cerevisiae by microarray hybridization. Mol. Biol. Cell 9, 3273–3297 (1998).

41. Wagner, F., Yan, Y. & Yanai, I. K-nearest neighbor smoothing for high-throughput single-cell RNA-Seq data. bioRxiv (2018). doi:10.1101/217737

42. Ferrezuelo, F., Colomina, N., Futcher, B. & Aldea, M. The transcriptional network activated by Cln3 cyclin at the G1-to-S transition of the yeast cell cycle. Genome Biol. 11, R67 (2010).

43. Liu, Z. et al. Reconstructing cell cycle pseudo time-series via single-cell transcriptome data. Nat. Commun. 8, 22 (2017).

44. Rustici, G. et al. Periodic gene expression program of the fission yeast cell cycle. Nat. Genet. 36, 809–817 (2004).

45. Dirick, L., Böhm, T. & Nasmyth, K. Roles and regulation of Cln-Cdc28 kinases at the start of the cell cycle of Saccharomyces cerevisiae. EMBO J. 14, 4803–4813 (1995).

46. Stuart, D. & Wittenberg, C. CLN3, not positive feedback, determines the timing of CLN2 transcription in cycling cells. Genes Dev. 9, 2780–2794 (1995).

47. Tyers, M., Tokiwa, G. & Futcher, B. Comparison of the Saccharomyces cerevisiae G1 cyclins: Cln3 may be an upstream activator of Cln1, Cln2 and other cyclins. EMBO J. 12, 1955–1968 (1993).

48. Wijnen, H., Landman, A. & Futcher, B. The G1 Cyclin Cln3 Promotes Cell Cycle Entry via the Transcription Factor Swi6. Mol. Cell. Biol. 22, 4402–4418 (2002).

49. Hartwell, L. H., Culotti, J., Pringle, J. & Brian Reid. Genetic control of the cell division cycle in yeast. Science 183, 46–51 (1974).

50. de Bruin, R. A. M., McDonald, W. H., Kalashnikova, T. I., Yates III, J. & Wittenberg, C. Cln3 Activates G1-Specific Transcription via Phosphorylation of the SBF Bound Repressor Whi5. Cell 117, 887–898 (2004).

51. Schwob, E. & Nasmyth, K. CLB5 and CLB6, a new pair of B cyclins involved in DNA replication in Saccharomyces cerevisiae. Genes Dev. 7, 1160–1175 (1993).

52. Lim, H. H., Goh, P.-Y. & Surana, U. Cdc20 is essential for the cyclosome-mediated proteolysis of both Pds1 and Clb2 during M phase in budding yeast. Curr. Biol. 8, 231–237 (1998).

53. Prinz Susanne, A. A. Dual control of mitotic exit. Nature 402, 133 (1999).

54. Shirayama, M., Tóth, A., Gálová, M. & Nasmyth, K. APCCdc20 promotes exit from mitosis by destroying the anaphase inhibitor Pds1 and cyclin Clb5. Nature 402, 203 (1999).

55. Visintin, R. et al. The phosphatase Cdc14 triggers mitotic exit by reversal of Cdk-dependent phosphorylation. Mol. Cell 2, 709–718 (1998).

56. Skotheim, J. M., Di Talia, S., Siggia, E. D. & Cross, F. R. Positive feedback of G1 cyclins ensures coherent cell cycle entry. Nature 454, 291–296 (2008).

57. Johnston, G. C., Pringle, J. R. & Hartwell, L. H. Coordination of growth with cell division in the yeast Saccharomyces cerevisiae. Exp. Cell Res. 105, 79–98 (1977).

58. Bähler, J. Cell-cycle control of gene expression in budding and fission yeast. Annu. Rev. Genet. 39, 69–94 (2005).

59. Shaltiel, I. A., Krenning, L., Bruinsma, W. & Medema, R. H. The same, only different – DNA damage checkpoints and their reversal throughout the cell cycle. J. Cell Sci. 128, 607–620 (2015).

60. Branzei, D. & Foiani, M. Regulation of DNA repair throughout the cell cycle. Nat. Rev. Mol. Cell Biol. 9, 297–308 (2008).

61. Hustedt, N. & Durocher, D. The control of DNA repair by the cell cycle. Nat. Cell Biol. 19, 1–9 (2017).

62. Murray, J. M. & Carr, A. M. Integrating DNA damage repair with the cell cycle. Curr. Opin. Cell Biol. 52, 120–125 (2018).

63. Roy, D. et al. Interplay between cancer cell cycle and metabolism: Challenges, targets and therapeutic opportunities. Biomed. Pharmacother. 89, 288–296 (2017).

64. Kaplon, J., van Dam, L. & Peeper, D. Two-way communication between the metabolic and cell cycle machineries: the molecular basis. Cell Cycle 14, 2022–2032 (2015).

65. Kalucka, J. et al. Metabolic control of the cell cycle. Cell Cycle 14, 3379–3388 (2015).

66. Huang, C. Y. & Ferrell, J. E. Ultrasensitivity in the mitogen-activated protein kinase cascade. Proc. Natl. Acad. Sci. 93, 10078–10083 (1996).

67. Steinway, S. N. et al. Network modeling of TGFβ signaling in hepatocellular carcinoma epithelial-to-mesenchymal transition reveals joint sonic Hedgehog and Wnt pathway activation. Cancer Res. 74, 5963–5977 (2014).

68. Venables, W. N. & Ripley, B. D. Modern Applied Statistics with S. (Springer, 2002).

